# Phylogeographic clustering of *Salmonella enterica* serovar Mississippi in the Southeastern United States indicates regional transmission pathways

**DOI:** 10.1101/2025.03.26.645529

**Authors:** Mel H. Yoshimoto, Lauren K. Hudson, Harleen K. Chaggar, Katie N. Garman, John R. Dunn, Agricola Odoi, Thomas G. Denes

## Abstract

*Salmonella enterica* subspecies *enterica* serovar Mississippi (*S*. Mississippi) is a polyphyletic serovar endemic in Australia, New Zealand, the United Kingdom, and the United States (USA). From 2018 to 2024, it was the 13^th^ most frequently reported clinical *Salmonella* serovar in the USA. Its incidence in the USA is geographically focused on the Southeast. From 2018 to 2024, 78% of all *S*. Mississippi cases in the USA occurred in Southeastern states. The objective of this study was to determine the phylogeographic patterns of clinical *S*. Mississippi in the Southeastern USA using clinical *S*. Mississippi isolate sequencing data and metadata from ten state public health laboratories in the region. The resulting phylogeny, based on core single nucleotide polymorphism (SNP) differences, showed that *S*. Mississippi has five major clades (Ai, Aii, Bi, Bii, C), four of which are consistent with the results of previous studies. Clade Ai, which comprised 99% of study isolates, was systematically divided into seven subclades. County-level mapping of the clade Ai isolates revealed distinct geographical distributions at the clade and subclade levels. For example, subclade Ai1 was predominantly identified along the East Coast, while subclade Ai3 was primarily found in Western Tennessee. Simple linear regression showed a significant positive association (p < 0.01) between isolate-to-isolate genomic distance (SNP differences) and county-to-county geographic distance (km) at the clade and subclade levels. These findings provide insight into the transmission of *S*. Mississippi and may inform public health hypothesis generation.

**IMPORTANCE:** This study enhances understanding of *S*. Mississippi transmission pathways and reveals localized phylogeographic clustering within the Southeastern USA. Classification of this serovar with consideration to geographical clustering may improve surveillance, investigation, and control efforts. Geographical clustering of this serovar within this region suggests local or regional transmission pathways and enzootic reservoirs, which could be related to environmental or regional food exposures. Identifying possible environmental sources, enzootic reservoirs, or climate factors contributing to human infection with this serovar within the Southeastern USA could lead to prevention recommendations. Knowledge of exposure patterns for *S*. Mississippi may facilitate development and implementation of targeted control strategies and interventions locally that can limit further exposure and prevent illness.

## INTRODUCTION

*Salmonella enterica* subspecies *enterica* serovar Mississippi (*S*. Mississippi) was the 13th most frequently reported clinical *Salmonella* serovar in the United States (USA) according to Report Data from 2018 to 2024 obtained from the BEAM Dashboard (1). *S*. Mississippi is a nontyphoidal serovar but has been found to encode the *Salmonella* cytolethal distending toxin (S-CDT). S-CDT is a genotoxin that may cause invasive systemic disease, persistent asymptomatic bacteremia, DNA damage, and other forms of long-term sequelae (2, 3). Rare but recorded sequelae caused by *S*. Mississippi include renal abscess in an immunocompetent patient (4) and transplacental transmission resulting in a spontaneous miscarriage at 18 weeks gestation (5).

Existing literature suggest that *S*. Mississippi is a polyphyletic serovar endemic in Australia, New Zealand, the United Kingdom, and the USA (6-8). The serovar is considered endemic to Tasmania, with wide genetic diversity and persistence in the environment, with a broad range of host reservoirs that persist over time (9). Previous phylogenetic analysis by Cheng et al. compared whole-genome sequence (WGS) data of *S*. Mississippi isolates with other *S. enterica* serovars. They identified two primary clades: clade A, within *S*. enterica and clade B, within section Typhi of *S*. enterica. Their results suggest that clades of *S*. Mississippi evolved from different ancestors, and that isolates cluster geographically. Clade A isolates within section Typhi were primarily from the USA (subclade Ai) and Australia (subclade Aii). Clade B isolates were primarily from the United Kingdom (6). The number of polyphyletic serovars known to us continues to increase as new *Salmonella* isolates continue to be sequenced (6).

*S*. Mississippi has historically been geographically focused within the Southeastern USA (10) and the Gulf Coast states (11) and has been a frequently reported serovar since 2010. Incidence of *S*. Mississippi is highest along the Mississippi River and the Mississippi-Louisiana coast (10, 12). From 2018 to 2024, 77.7% of all *S*. Mississippi cases in the USA occurred in the Southeast (Alabama [AL], Arkansas [AR], Florida [FL], Georgia [GA], Kentucky [KY], Louisiana [LA], Mississippi [MS], North Carolina [NC], South Carolina [SC], Tennessee [TN], and Virginia [VA]) (1). The highest incidence rates of culture-confirmed human *S*. Mississippi infections (per 100,000 population) in 2016 were in MS (4.2), LA (2.1), and NC (1.0), followed by AL (0.8), TN (0.6), AR (0.5), and GA (0.3) (10). KY (0.1), SC (0.1), VA (0.1), and FL (0.02) had lower incidence compared with other Southeastern states (10). Geographical restriction or distribution may indicate association with local habits or food products, persistence in, or adaptation to particular natural environments, and/or animal reservoirs within a specific habitat (9, 11, 13, 14).

*S*. Mississippi may have higher transmission rates from animal or environmental sources than food (8, 15, 16). According to National Outbreak Reporting System (NORS) data obtained from the BEAM Dashboard, there have been only four outbreaks caused by *S*. Mississippi with confirmed etiology (and two with suspected etiology) (17). Of those, only one lists a food vehicle: a 2022 multistate outbreak caused by tomatoes resulting in 102 illnesses and 23 hospitalizations. Among the top 20 nontyphoidal *Salmonella* serovars in the USA, *S*. Mississippi had the lowest foodborne relatedness (FBR) measure (0.01) (18). This measure indicates a low association with foodborne transmission.

Previous *S*. Mississippi studies show evidence of human infection from wildlife. This serovar has been associated with or isolated from zoonotic reservoirs, including wild animal (7, 12), avian (8, 12), amphibian, reptile (12), and equine sources (6, 12)). Other environmental sources identified include untreated drinking water (7, 8) and untreated recreational water (2018 outbreak in Alabama associated with a lake/reservoir) (17). Untreated freshwater reservoirs and watersheds, such as the Mississippi River, may serve as natural reservoirs for *S*. Mississippi (19).

Additionally, *S*. Mississippi exhibits high seasonality, with summer peaks exceeding other *Salmonella enterica* serovars (7, 8, 11). This temporal increase could be related to increased presence of animal host(s) (e.g., reptiles, amphibians), increased exposure to a source (e.g., recreational water), or increased pathogen levels in a continually present source due to increasing favorable environmental conditions (11, 20). Identifying control strategies for this serovar is difficult due to its high genetic diversity and wide range of animal and environmental sources (7-9, 12).

The purpose of this study was to better understand spatial patterns of *S*. Mississippi in the Southeastern USA and describe the relationship between genomic distance and geographical distance. We determined the phylogeny of *S*. Mississippi clinical isolates from ten Southeastern USA states and performed county-level exploratory spatial data analysis (ESDA) of sequenced clinical *S*. Mississippi isolates, providing increased resolution on how this serovar is distributed within the region. Additionally, we assessed the association between genomic and geographic distance using simple linear regression to develop our knowledge on the county-level population ecology of this serovar, informing future outbreak investigations.

## RESULTS

### Phylogeny consists of five clades

Phylogenetic analyses of the study isolates (n = 2,797) and reference genomes (n = 56) revealed five primary clades: Ai, Aii, Bi, Bii, and C (**Figure 1A**). This shows that *S*. Mississippi is a polyphyletic serovar, consistent with the results of Cheng et al. (6). Four of the identified clades (Ai, Aii, Bi, and Bii) correspond to those identified by Cheng, et al. However, an additional fifth clade (C) was identified that was not observed by Cheng and colleagues. Notably, clade C only contained three isolates. Clades Ai and Aii are more closely related, with clade C falling between A clades and B clades. Clade Aii only contained one study isolate (from NC) and the reference genomes in this clade (n = 19) were primarily from New Zealand (n = 9) and Australia (n = 9). Clade Bi contained 22 study isolates (from all included states, except AL and AR). The reference genomes in this clade (n = 16) were primarily from the USA (n = 15). Clade Bii contained no study isolates, and all the reference genomes contained in this clade (n = 7) were from the United Kingdom. Clade C only contained three study isolates, all from VA (6).

**Figure 1.**
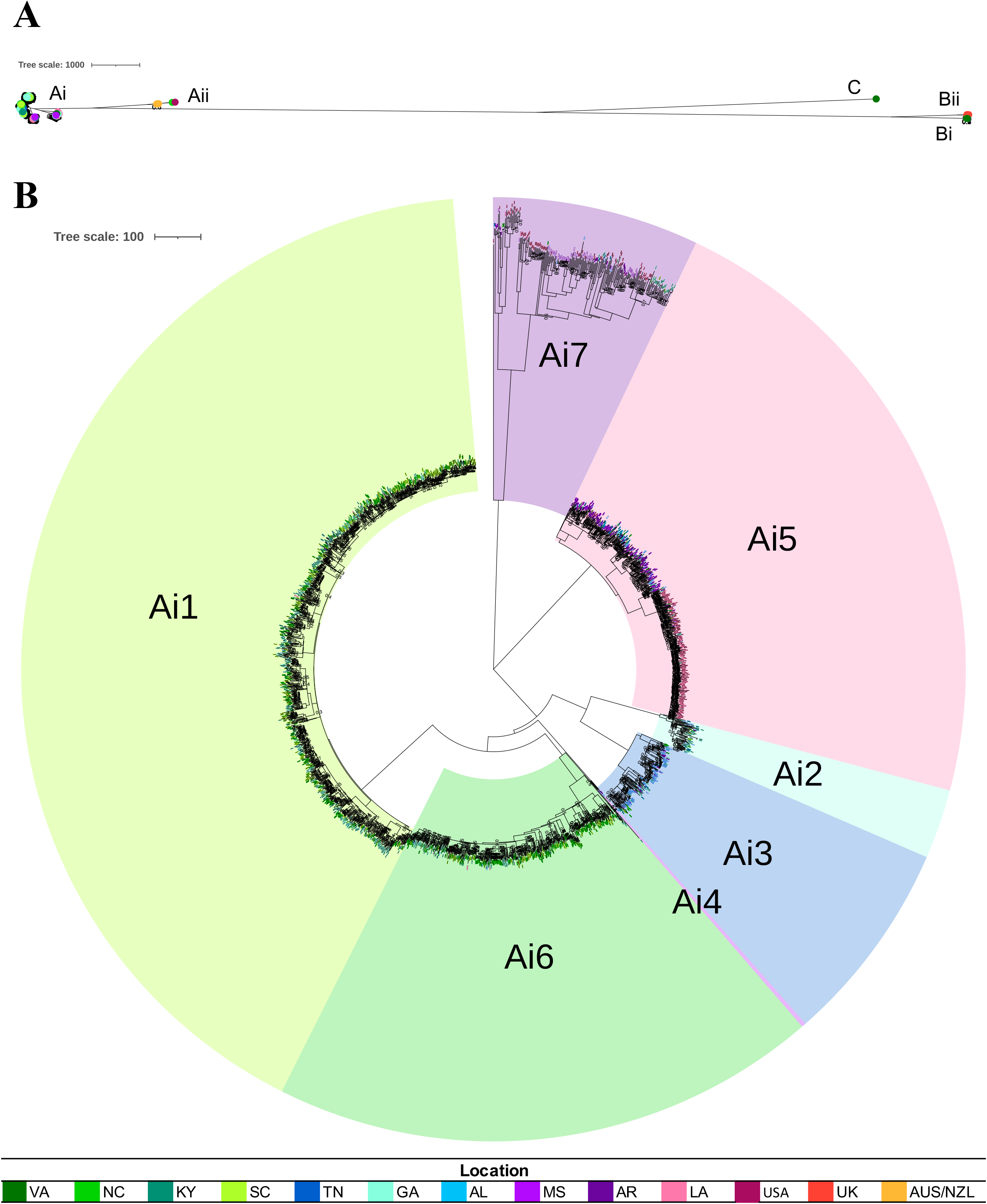
Neighbor joining phylogenetic trees of all isolates and of clade Ai with subclades. Both trees were constructed in MEGAX (46, 58) using the neighbor-joining method (59) and visualized and annotated using iTOL (48). The trees are drawn to scale, with branch lengths representing the number of base differences at core SNP positions. The evolutionary distances were computed using the number of differences method (60). All ambiguous positions were removed for each sequence pair (pairwise deletion option). Isolate locations are indicated by colored shapes or text at tree tips (see legend). (A) Phylogeny of all isolates. This analysis involved 2853 isolates and a total of 55,550 core SNP positions. The proportion of replicate trees in which the associated taxa clustered together in the bootstrap test (50 replicates) are shown next to the branches if ≤0.7 (61). (B) Phylogeny of clade Ai. This analysis involved 2775 isolates and a total of 40,891 core SNP positions. The proportion of replicate trees in which the associated taxa clustered together in the bootstrap test (100 replicates) are shown next to the branches if ≤0.7 (61). Subclades are indicated.

Clade Ai contained approximately 99% of the study isolates (n = 2,761) and reference genomes from the USA (n = 14). This is consistent with the results of Cheng, et al. (6), who found that clade Ai isolates were predominantly from the USA. Further analyses focused on this clade. Clade Ai was systematically divided, resulting in seven Ai subclades (**Figure 1B**).

Subclades Ai1 and Ai6 isolates were primarily from VA, NC, SC, and GA. Subclade Ai2 isolates were primarily from GA, with a small but notable portion of isolates from TN and AL. Subclade Ai3 isolates were primarily from TN, with several isolates from VA and KY. Subclade Ai4 isolates were from TN, MS, and GA. Close to half of subclade Ai5 isolates were from LA, with the other half predominantly from MS followed by AL and TN. Subclade Ai7 isolates were primarily from AR and LA, followed by GA.

### County-level geospatial patterns

The geographical distribution of *S*. Mississippi isolates per county population by clade shows that clade Ai was clustered in two geographical regions of the Southeastern USA (**Figure 2A**): (i) One high risk region runs from West TN to Southeast LA, covering Northeast to Southwest MS; follows the Mississippi River (on the MS side) except for a portion of Northwest MS and (ii) the other high risk region runs throughout states bordering the East Coast, from the VA-NC border, throughout most counties in SC, and along the SC-GA border. The highest numbers of clinical *S*. Mississippi isolates per county population was observed in South-Central NC, Central SC, and Southeastern GA.

**Figure 2.**
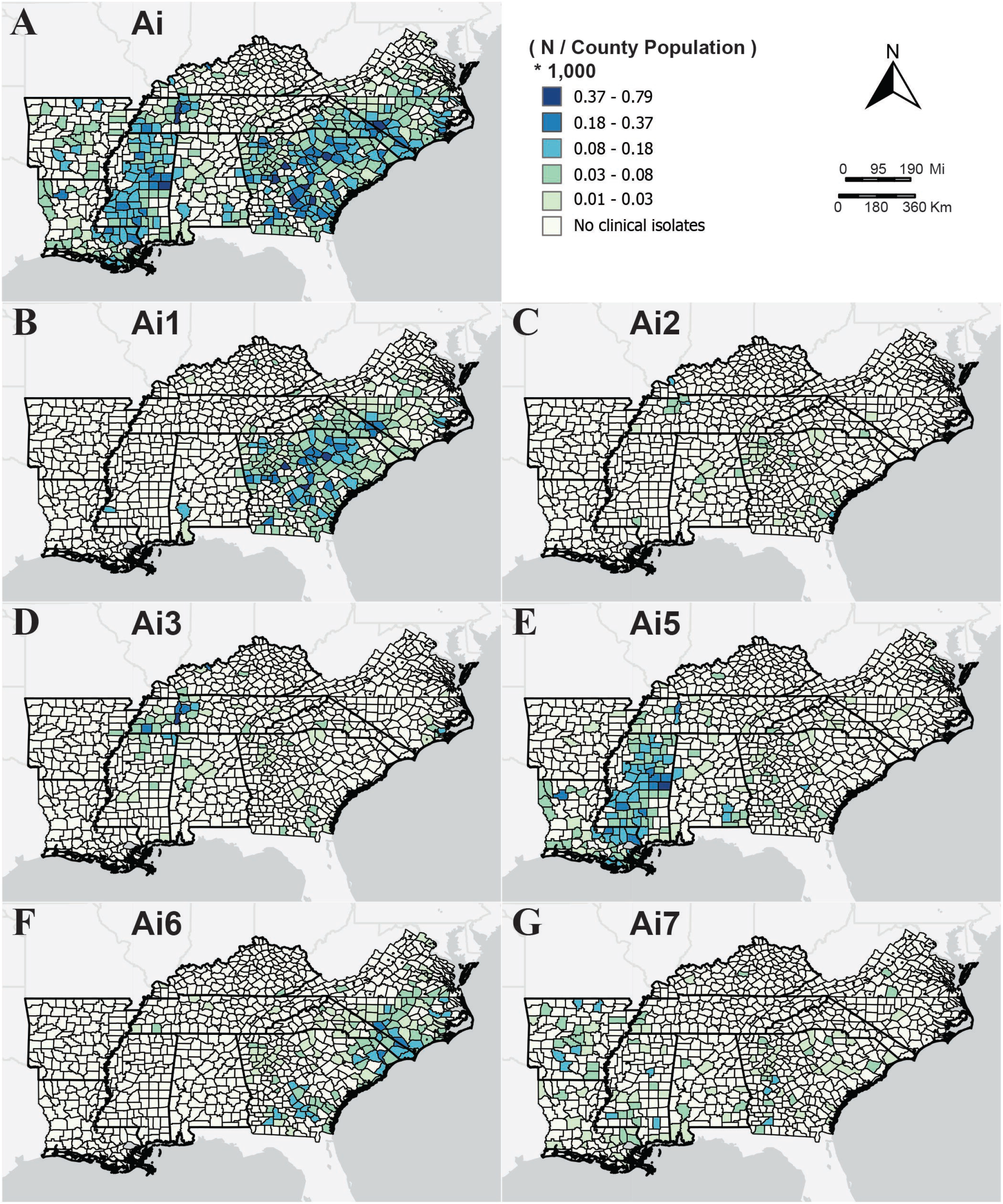
County-level risk of *S*. Mississippi clinical isolates by Clade. Counties are shaded by number of *S*. Mississippi isolates per county population (see legend at top right) for (A) all of clade Ai and (B-H) subclades Ai1 through Ai7, excluding subclade Ai4.

Close examination of the geographical distribution of subclades of clade Ai reveals that subclades Ai1, Ai3, Ai5, and Ai6 had the strongest evidence of spatial clustering in the Southeastern USA (**Figures 2B-2G**). There is overlap in the geographic distribution of subclades Ai1 and Ai6. Subclade Ai1 (**Figure 2B**) was predominantly distributed along the East Coast from Southern VA through SC and most of GA, with other occurrences in Clarke and Baldwin counties in AL and Jefferson County in MS. Subclade Ai6 (**Figure 2F**) exhibited a strong presence along the Eastern Seaboard from Southeast VA to Southeast GA, with inland distribution in NC, SC, and GA, particularly notable in NC. This subclade also had a sparse and scattered distribution throughout TN, KY, and inland VA.

There is also overlap in the geographic distribution of subclades Ai3 and Ai5. Subclade Ai3 (**Figure 2D**) was primarily clustered in Western TN and Alcorn and Tishomingo counties in MS, Hancock County in KY, and Pamlico County in NC. Subclade Ai5 (**Figure 2E**) was observed in West-Central TN through Northeast MS to Southeast LA (mostly in the Southern Deltas, but also the Mississippi Alluvial Plain), with a moderate presence in Northwest LA counties and Southeastern AL.

In comparison, the distributions of subclades Ai2 and Ai7, in addition to being less common, were more sporadic spatially. Subclade Ai2 (**Figure 2C**) was sparsely distributed throughout KY, TN, NC, MS, AL, and GA, with the most notable presence in Northern GA and West-Central TN. The highest risk was observed in Livingston County in KY (the only county in KY for this subclade), Houston County in TN, and McIntosh County in GA. The distribution of subclade Ai7 (**Figure 2G**) was random throughout the region, similar to subclades Ai2 and Ai4, with exceptions in Southwest MS and Central AR. The highest risks of these isolates were observed in AR, MS, and GA. In AR and GA, the highest risk was in more densely populated counties, while in MS, it was more common in rural counties.

Subclade Ai4 (**Figure S1A**) contained fewer than five total isolates and was limited to Tate and Yalobusha counties in MS and Fayette County in TN.

### Association between spatial and genomic distances

Pairwise SNP differences were skewed to the left because we are looking at isolates from the same serovar (**Figure 3**). The Box-Cox method suggested the square root transformation on the response variable, geographical distance (km). The association between the square root of pairwise geographical distance in km (response/dependent variable) and pairwise genomic distance in SNP differences (predictor/independent variable) was measured using simple linear regression (SLR) for individual subclades within clade Ai and for clade Ai as a whole.

**Figure 3.**
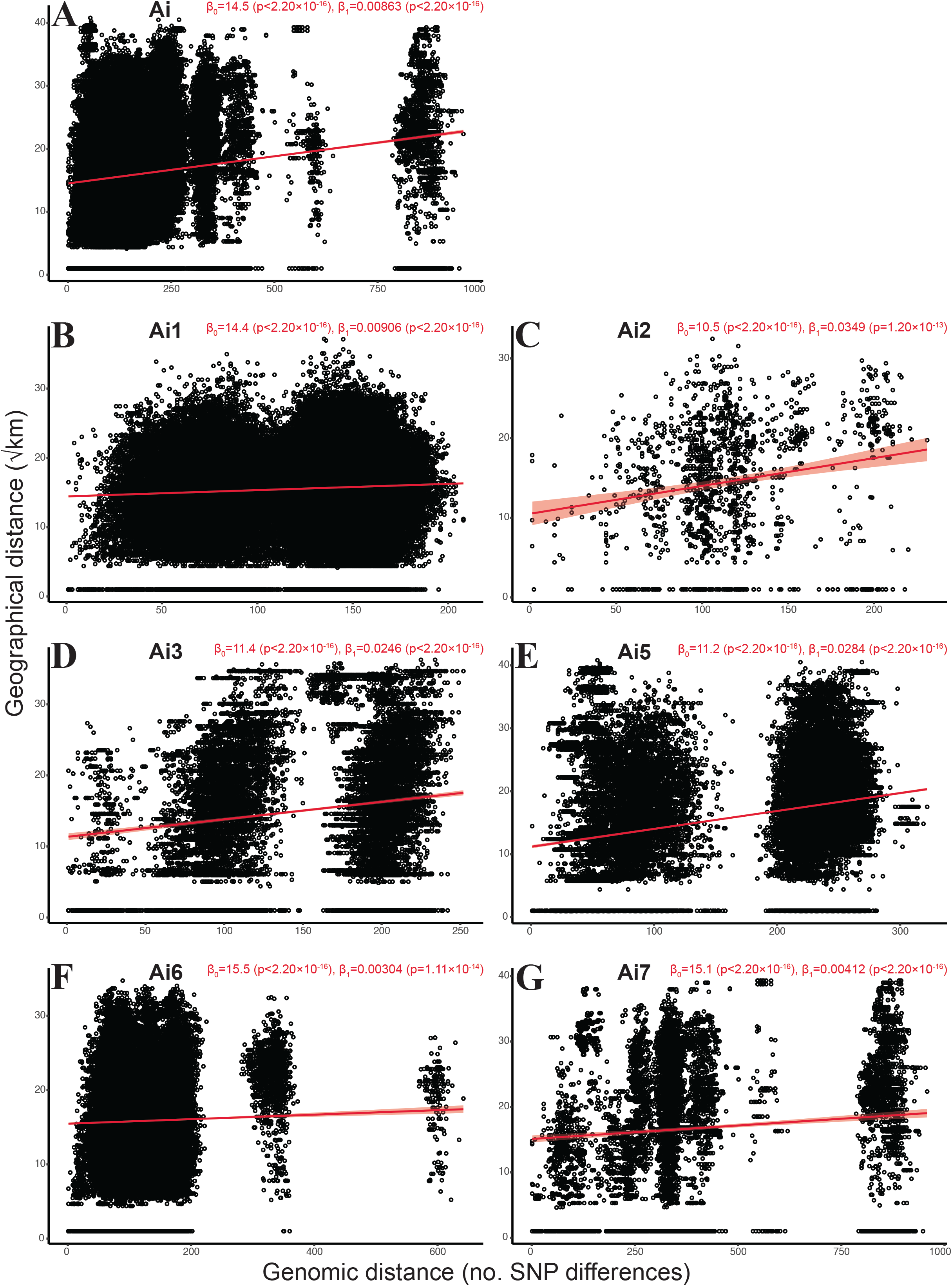
Scatterplots of genomic distance vs. geographical distance between *S*. Mississippi isolates. Scatterplots showing genomic distance (no. SNP differences) vs. geographical distance (km, square root transformed) for (A) all of clade Ai and (B-H) subclades Ai1 through Ai7, excluding subclade Ai4. For each, the red line and light red shading represent the regression line and 99% confidence intervals of the simple linear model. The y-intercept (β_0_), slope (β_1_), and their associated p-values are in red at the top right of each panel. F statistics and model p-values are provided in **Table S3**.

All subclades, except subclade Ai4 (**Figure S1B**), showed significant positive associations between genomic and geographical distances (**Figure 3**). The lack of significant association for subclade Ai4 might be attributed to the small sample size (n = 6). There was evidence of significant association between genomic and geographic distance even when all clade Ai subclades were combined. When comparing the clade Ai data containing only within-group (within each subclade) comparisons to the clade Ai data with between-group (between-subclades) comparisons, the data with between-group comparisons had a slightly larger positive magnitude. SLRs for both datasets had a very small p-value (p < 0.001).

## DISCUSSION

The objective of this study was to determine the phylogeographic patterns of clinical *Salmonella enterica* subspecies *enterica* serovar Mississippi (*S*. Mississippi) in the Southeastern USA using a large dataset of clinical isolates from this region (n = 2,739). We found that *S*. Mississippi shows distinct geographical associations at the clade and subclade level. The geographical clustering of this serovar within the Southeastern USA suggests local or regional transmission pathways and reservoirs, which could be related to environmental exposures or regional food exposures (20). This pattern differs from other serovars that show relatively uniform geographic distribution, such as Typhimurium or Enteritidis, which indicates uniform sources that are widely dispersed (e.g., nationally distributed food products) or a balanced mix of multiple sources of exposure (11). Surveillance data showing that *S*. Mississippi appears to be more associated with sporadically acquired infections than outbreak-associated (only 0.83% clinical isolates from 2018-2024) (1) further supports that exposures are more likely to be environmental (21). Some potential environmental or ecological factors contributing to the observed spatial distribution in this study are principal aquifers (22), watersheds (23), agriculture (24), bird migration patterns (7), amphibian breeding (9, 20), population densities (24), temperature (25, 26), or ecosystem categorization (e.g., wetland type) (27).

Supporting our observation of geographical clustering, we found statistically significant positive association between genomic and geographical distances at both the clade and subclade levels (except for subclade Ai4, where N < 10). Thus, the number of SNP differences between two clinical isolates of *S*. Mississippi could possibly be used to predict the geographic location of exposures that resulted in the isolate-associated illnesses. However, further research is needed to understand the mechanism(s) of this positive relationship and how it varies between subclades.

As genomic-based methods are being developed and refined for public health implementation, there is a need to determine the optimal level of genomic resolution (e.g., genomic distance thresholds) for subtyping of specific pathogen species or serovars for surveillance applications (e.g., cluster detection) (28). This is also needed for studies looking at “functional clades” similar to the current study (at what level of relatedness can geospatial and other trends be seen?). Cheng, et al. found that this serovar has associations with differing countries at the clade level: Ai with the USA, Aii with Australia, and Bi and Bii with the United Kingdom (6). The findings here extend the work by Cheng, et al. (6) on the phylogeny of this serovar, with a specific focus on the population structure of isolates from the Southeastern USA, which are predominantly part of clade Ai. The subclade-level subtyping employed in the current study shows that this finer level of resolution reveals functionally relevant subclades where distinct geographical clustering evident, which could be used to inform regional and state level public health response. For example, the phylogeographic distributions identified in the current study could be used by public health professionals, along with epidemiological data, to determine the potential geographic origin of a contamination or exposure event based what clade an isolate belongs to (6, 29, 30).

County-level spatial analysis of clinical isolates and measuring the association between genomic and geographical distances for this serovar will allow for future exploration and identification of localized environmental risks. Environmental sampling to isolate and characterize environmental *S*. Mississippi isolates, similar to the nationwide genomic atlas of soil-dwelling *Listeria* developed by Liao, et al. (31), could assist in identifying risk factors and natural reservoirs. Additionally, as temporal differences were not taken into account in the current study, it would be helpful to assess temporal incidence and climate factors (e.g., rainfall, temperature) or look at environmental factors that have a spatial pattern (e.g., landcover indices for water) to determine any correlations with incidence of human disease.

Identifying control strategies for polyphyletic serovars like *S*. Mississippi can prove difficult, as they are typically associated with a wide range of animal and environmental sources (7-9, 12). For other polyphyletic serovars, research has shown that different lineages can be associated with different hosts (6, 32, 33). A better understanding of sources of exposure will facilitate the development and implementation of targeted education and control strategies and interventions that limit further exposure and prevent illness. Additionally, knowledge of transmission pathways can aid in the design of questionnaires to gather relevant risk factor and exposure data and earlier identification of outbreak sources, leading to a more robust public health response.

The transition to WGS for surveillance of enteric pathogens has created an opportunity to integrate epidemiological and high-resolution genomic data to better understand pathogen transmission dynamics, as was done in the current study and can be applied to other *Salmonella* serovars and foodborne pathogens. This approach can also be used to elucidate sources or reservoirs with a more targeted subgroup of pathogens, such as reoccurring, emerging, or persisting (REP) strains, a new classification used by the CDC to track strains that cause illnesses over longer periods of time than an acute outbreak (34).

One limitation of this study is the use of isolates obtained through case-based passive surveillance, which is susceptible to underreporting or other biases. Also, because isolate data was obtained from different state public health departments, there may be differences between their surveillance, isolate submission, characterization, or data collection practices (e.g., for county of isolation, some states may use county the county where the isolate was collected and others may use the county of residence of the case). Additionally, exact exposure information was not available, so, in some cases, county of isolation may not be where the exposure occurred.

In this study, we examined the phylogeography of *S*. Mississippi in the Southeastern USA, a region where this serovar is geographically focused. We found a significant (p < 0.01) positive association between isolate-to-isolate SNP differences and the square root of county-to-county geographic distance at both the clade and subclade levels. This information could be leveraged by public health professionals to use genomic distances to predict geographic locations of exposures that result in illnesses. Additionally, we found that this serovar shows distinct phylogeographic distributions at both the clade and subclade levels. These findings provide insight into the transmission of *S*. Mississippi. The localized phylogeographic clustering within this region suggests local or regional transmission pathways and reservoirs, which could be related to environmental or regional food exposures. This information can be leveraged to further investigate and identify possible environmental sources, enzootic reservoirs, or climate factors contributing to human infection with this serovar within the Southeastern USA.

## MATERIALS AND METHODS

### Isolate information

A multistate exploration of sequenced *Salmonella* Mississippi clinical isolates was conducted by collaborating with state public health laboratories (SPHL) in the Southeastern USA (AL, AR, GA, KY, LA, MS, NC, SC, TN, and VA). The following information was requested from each SPHL on all sequenced *S*. Mississippi clinical isolates received up to mid-2023: isolate identifiers (PNUSA, SRR, or SAMN), source county, and year of isolation (**Table 1, Table S1**). NCBI E-utilities were used to retrieve SRR identifiers for isolates where only PNUSA was provided. Fasterq-dump (v2.11.2, part of SRA Toolkit) (35) was used to retrieve raw reads FASTA files (n=2,797) from the NCBI SRA database. The raw reads were trimmed using Trimmomatic (v0.39, Phred: 33, Leading: 3, Trailing: 3, Sliding window: 4:15, Min len: 36 (36). Quality control was performed using FastQC (v0.11.9) (37) and MultiQC (v1.11) (38) Trimmed reads were assembled into contigs using SPAdes (v3.13.1, (39, 40), with the careful option). Contigs with a length of <1 kb or coverage <5 × were removed from assemblies (41). QUAST (v5.0.2 (42)) was used to generate assembly statistics.

**Table 1.** Count of sequenced *S. Mississippi* clinical isolates per state, including no. isolates with known source county.

| State | No. isolates |  |  |  |
| --- | --- | --- | --- | --- |
|  | Total | Raw reads available | County known | Raw reads available and county known |
| Alabama | 61 | 57 | 61 | 57 |
| Arkansas | 75 | 58 | 73 | 58 |
| Georgia | 688 | 673 | 648 | 634 |
| Kentucky | 15 | 15 | 15 | 15 |
| Louisiana | 388 | 388 | 387 | 387 |
| Mississippi | 229 | 229 | 229 | 229 |
| North Carolina | 627 | 605 | 611 | 593 |
| South Carolina | 465 | 465 | 459 | 459 |
| Tennessee | 194 | 191 | 194 | 191 |
| Virginia | 109 | 66 | 85 | 66 |
| Other/unk. | 52 | 50 | 0 | 0 |
| **Total** | **2,903** | **2,797** | **2,762** | **2,689** |

### Phylogenetic analyses

A total of 56 reference genomes were included in the phylogenetic analysis (**Table 2, Table S2**). Nineteen complete assemblies of clinical *S*. Mississippi isolates were downloaded from NCBI. Thirty-seven reference genomes from Cheng et al. (6) were included in the analysis to place the current study isolates into the context of already established clades. Reference-free SNP detection was performed using KSNP (v3.1, k = 19 (43-45)) with the assembled isolates (n = 2,797) and reference genomes retrieved from NCBI (n=56). The resulting core SNP matrix fasta file was processed in MEGA (v10.2.6 (46)) to construct a neighbor-joining phylogenetic tree with 50 bootstrap replicates. The tree was further visualized and annotated in iTOL (47, 48) The initial phylogenetic tree, the following assembly statistics (percent G+C [51.9-52.5%], total length [4.5-5.0 Mb], number of contigs [22-217], and estimated coverage [20-275 ×]), and previous studies (41) were used to develop the inclusion criteria: 51.9-52.4% G+C, 4.4-5 Mb length, ≤200 contigs, estimated coverage ≥10 ×. Exclusions (n = 10) were made accordingly (13).

**Table 2.** Count of sequenced *S. Mississippi* clinical isolates per clade and subclade.

| Clade/Subclade | Study | Reference | Total |
| --- | --- | --- | --- |
| **Ai** | **2,771** | **14** | **2,785** |
| Ai1 | 1,155 | 5 | 1,160 |
| Ai2 | 67 | 0 | 67 |
| Ai3 | 197 | 0 | 197 |
| Ai4 | 4 | 0 | 4 |
| Ai5 | 619 | 2 | 621 |
| Ai6 | 522 | 5 | 527 |
| Ai7 | 197 | 2 | 199 |
| **Aii** | **1** | **19** | **20** |
| **Bi** | **22** | **16** | **38** |
| **Bii** | **0** | **7** | **7** |
| **C** | **3** | **0** | **3** |
| **Total** | **2,797** | **56** | **2,853** |

The majority of the *S*. Mississippi clinical isolates were within clade Ai (n=2,761). KSNP was re-run with only clade Ai isolates for increased resolution of genomic distance between the clinical isolates. The resulting core SNP matrix fasta file was used to create a phylogenetic tree as previously described. FastBAPS (v1.0.8, initial hierarchy algorithm (49)) was used to systematically determine subclades within clade Ai, and KSNP was run separately with each of the seven resulting clade Ai subclades (Ai1-Ai7). The resulting core SNP matrix fasta files were used to create phylogenetic trees as previously described.

### Spatial analysis and mapping

County-level risk of *S*. Mississippi (n = 2,667) was mapped using ArcGIS Pro (v 3.3 (50)) and TIGER/Line shapefiles containing population data from the United States Census Bureau FTP Archive (51). A total of 72 isolates could not be included in the spatial analysis due to missing county of isolation; these isolates do not have the geographical linkage required to perform a spatial analysis. County-level risk was computed by dividing the total number of sequenced isolates from each county by the county population and applying a population multiplier of 1000. The county-level spatial distribution of the isolates within clade Ai was mapped. Individual maps for each subclade were generated and their spatial patterns visually compared.

### Descriptive statistical analysis

Data processing and statistical analysis were conducted using R (v4.3.2; (52)) in RStudio (v2023.12.1+402; (53)). The tidyverse library (54) was used to clean, parse, and analyze data for spatial and genomic distance for clade Ai. Spatial distance was defined as the pairwise distance between the geographic centroids of counties of isolation for isolates in the study population. All combinations of pairwise county-to-county distances were retrieved from the National Bureau of Economic Research. These distances are great-circle distances, calculated using the Haversine formula (55). Pairwise distances for county and SNP data were cleaned, with geographical data restricted to counties within the states of AL, AR, GA, KY, LA, MS, NC, SC, TN, and VA for isolates with a known county of origin. Isolates with no known county of origin were excluded. County distances were converted from miles to kilometers. Genomic distance was defined as the pairwise SNP difference between individual isolates; pairwise SNP differences were calculated using MEGA and exported in CSV format.

Isolate identifiers (SRR IDs) were parsed together with their respective county of isolation (5-digit FIPS code) to combine county and SNP difference data.

The psych library (psych::describe()) (56) was used to get a general idea of the variables’ distributions. Histograms of SNP difference and km distance were constructed using hist() and scatterplots were created using plot() + abline() to visually assess the distribution of the variables and how they might be correlated. The MASS library (MASS::bc()) (57) was used to transform the spatial distances using the Box-Cox method, making their distribution more normal. The Box-Cox method suggested the square root transformation on the response variable, geographical distance (km) (**Table S3**).

After transforming spatial distance based on the lambda provided by bc() (**Table S3**) and reassessing the distribution with hist(), lm() was used to perform simple linear regression (SLR) to measure the association between the square root of pairwise geographical distance in km (response/dependent variable) and pairwise genomic distance in SNP differences (predictor/independent variable) for the *S*. Mississippi clinical isolates. This process was repeated on individual data frames using SNP differences from individual KSNP runs for each of the seven subclades. A second data frame for clade Ai was constructed by parsing together individual subclade data frames using rbind(). Unlike the initial data frame for clade Ai, this data frame only contained within-group pairs.

## ACKNOWLEDGEMENTS

This study was supported by the Integrated Food Safety Centers of Excellence (Food Safety CoEs) through Epidemiology and Laboratory Capacity for Infectious Diseases (ELC) grant number NU50CK000528, multistate project S1077 “Enhancing Microbial Food Safety by Risk Analysis,” and by The University of Tennessee Institute of Agriculture AgResearch.

We would like to thank the following state health departments for providing the isolate information necessary for this project: Alabama Department of Public Health, Division of Infectious Diseases & Outbreaks; Arkansas Department of Health, Office of the Chief Science Officer; Georgia Department of Public Health, Georgia Public Health Laboratory; Kentucky Department for Public Health, Division of Laboratory Services; Louisiana Department of Health, Louisiana Office of Public Health; Mississippi State Department of Health, Mississippi Public Health Laboratory; North Carolina Department of Health and Human Services*, Division of Public Health, North Carolina State Laboratory of Public Health; South Carolina Department of Public Health, Advanced Molecular Detection Laboratory; Tennessee Department of Health, Foodborne and Enteric Diseases Program; and Virginia Department of Health, Division of Surveillance and Investigation. We would also like to thank Dr. Daleniece Higgins Jones for discussions supporting the project.

* The findings and conclusions in this publication are those of the author(s) and do not necessarily represent the views of the North Carolina Department of Health and Human Services, Division of Public Health.

## Author contributions

Conceptualization, TGD, MHY, LKH;

Data curation, MHY, LKH;

Formal analysis, LKH, MHY, AO;

Funding acquisition, TGD, KNG, JRD;

Investigation, LKH, MHY;

Methodology, LKH, MHY, AO

Project administration, TGD;

Resources, TGD, KNG, JRD;

Software LKH, HKC, MHY, AO;

Supervision, LKH, TGD;

Validation, MHY, AO;

Visualization, MHY, LKH;

Writing – original draft, MHY, LKH, TGD;

Writing – reviewing & editing, LKH, HKC, TGD, KNG, JRD, AO.

## Conflict of interest

The authors have declared no conflict of interest.

